# Reversing anterior insular cortex neuronal hypoexcitability attenuates compulsive behavior in juvenile rats

**DOI:** 10.1101/2021.10.28.466154

**Authors:** Kshitij S. Jadhav, Aurélien P. Bernheim, Léa Aeschlimann, Guylène Kirschmann, Isabelle Decosterd, Alexander F. Hoffman, Carl R. Lupica, Benjamin Boutrel

## Abstract

Development of self-regulatory competencies during adolescence is partially dependent on normative brain maturation. Here we report that juvenile rats as compared to adults exhibit impulsive and compulsive-like behavioral traits, the latter being associated with lower expression of mRNA levels of the immediate early gene zif268 in the anterior insula (AI). This observation suggests that deficits in AI function in juvenile rats could explain their immature pattern of interoceptive cue integration in rational decision-making and compulsive phenotype. In support of this, here we report hypoexcitability of juvenile layer-V pyramidal neurons in the AI, concomitant with reduced glutamatergic synaptic input to these cells. Chemogenetic activation of the AI attenuated the compulsive trait suggesting that delayed maturation of the AI results in suboptimal integration of sensory and cognitive information in adolescents and this contributes to inflexible behaviors in specific conditions of reward availability.

## Introduction

Compelling evidence demonstrates that heightened sensation seeking, together with risk taking and reckless behaviors, are a major cause of morbidity and mortality among teenagers ^1, 2^. Clinical evidence suggests that adolescents engage in dangerous activities despite knowing and understanding the risks involved ^3^, emphasizing that adolescents remain vulnerable to impulsiveness (i.e. acting prematurely without adequate forethought) due to incomplete development of executive cognitive functions (planning, abstract reasoning and response inhibition) ^4^. Therefore, adolescence is not only a transition phase from childhood to adulthood, but a normative process defined by the emergence of a sagacious mind shaped by the intricate influence of multiple experiences made of social pressure and adjustments in personal goals. Meanwhile, teenagers often display careless behaviors, particularly when acting recklessly is perceived as necessary for increased peer recognition ^3, 5, 6^. Thrill seeking in adolescence should not necessarily be perceived as a negative developmental trait, but one that can also be considered instrumental in shaping the adolescent brain to develop cognitive control through multiple experiences ^7^. As the underpinnings of adolescent risk-taking behaviors are poorly characterized, knowledge of the neurobiological mechanisms involved in this developmental stage is important to better define this transformative phase of brain circuitry.

Near the beginning of adolescence, the brain undergoes several structural and network reorganizations ^8, 9^. The discrepant trajectories of development characterizing the adolescent brain support several theories of increased adolescent risk-taking ^3, 10, 11, 12, 13, 14^; the core of all these hypotheses is the asynchronous development of neural systems underlying reward seeking and self-regulation ^15^. The limbic system matures earlier than the prefrontal cortex (PFC) and could be one reason for increased reward sensitivity resulting in increased sensation seeking ^16, 17, 18, 19^. It is argued that the delayed maturation of the PFC may represent increasing, but still incomplete, frontal control over behavior during adolescence ^20, 21^, gradually facilitating cognitive capacities for risk assessment ^14, 22, 23, 24, 25^. Whereas the mechanisms by which self-regulatory and affective brain networks interact remain unclear, converging evidence suggests that the insular cortex is critical for emotion regulation, cognitive control, and ultimately flexible behavior ^26, 27, 28, 29, 30^. Therefore, the tendency of adolescents to engage in risky behavior may also be due to their inability to engage harm avoidance circuitry including the anterior insular cortex (AIC) during decision making ^31^. Within this perspective, extensive review of the literature suggests that the key role of the AIC is in the integration of top-down cognitive predictions and bottom-up interoceptive signals for emotional awareness ^32^, contributing critically to the cognitive control network implicated in the coordination of thoughts and actions ^33^. In other words, the most adaptive behavioral response is predicated upon the integration and representation of all available cues, as well as the retrieval of representations based on past experiences in similar contexts. Hence, considering the linear pattern of grey matter maturation within the AIC during adolescence ^34^, it is not surprising that this brain area is considered critical for the development of self-regulatory competencies during adolescence. Acquiring a growing control over the integration of cognitive and affective information is critical for optimizing behavioral responses, specifically in the context of risky decision making ^28, 35^. However, a causal demonstration of this phenomenon is lacking.

In the present study, we first report a series of similar general coping strategies among juvenile and adult rats suggesting comparable capacities in the integration of cognitive and emotional information. However, in contrast to adults, adolescent rats showed higher impulsive and compulsive behaviors, the latter being associated with decreased mRNA expression of zif268 in the AIC. Then, using *in vitro* electrophysiology we describe lower excitability of layer 5 pyramidal neurons and smaller synaptic glutamatergic inputs to these cells in the AIC of juvenile rats. To determine the relevance of this AIC neuron hypoexcitability in adolescence, we chemogenetically activated AIC neurons in juvenile rats and find that this reduces compulsive behavior. Overall, our observations provide strong evidence that continued reward taking behavior in the face of aversive consequences depends, at least partially, on the decreased functional recruitment of AIC neurons in juvenile rats.

## Results

### Juvenile and adult rats display similar behavioral responses in absence of conflictual decision making

Since the AIC is considered an integrative hub that coordinates the recruitment of task- and context-relevant brain networks as a response to arousal ^29^, we conducted a series of behavioral experiments in adolescent and adult rats to assess their emotional coping strategies ^36^. First, juvenile (PND 40-50) and adult (PND 90-100) rats exhibited comparable locomotor activity in an open field (Figure 1A) and manifested equal preference for a novel environment (Figure 1B), suggesting similar emotional response and exploratory behavior to an unknown but neutral environment. We then challenged them in conditions of stress-induced arousal. In the passive avoidance paradigm, both groups similarly increased the latency to enter a dark compartment after previously experiencing mild electrical foot shocks in this environment (Figure 1C). In the contextual fear conditioning, juveniles exhibited a transiently enhanced freezing response after the first exposure to electrical foot shocks but with additional trial expressed similar freezing responses compared to adults (Figure 1D). Finally, an enhanced emotional response was found in juvenile rats exploring the elevated plus maze, reflected by reduced exploration of the open arms compared to adults (Figure 1E), despite similar levels of locomotion. Overall, exploratory behaviors in stressful environments only moderately differed between juvenile and adult rats.

**Figure 1.**
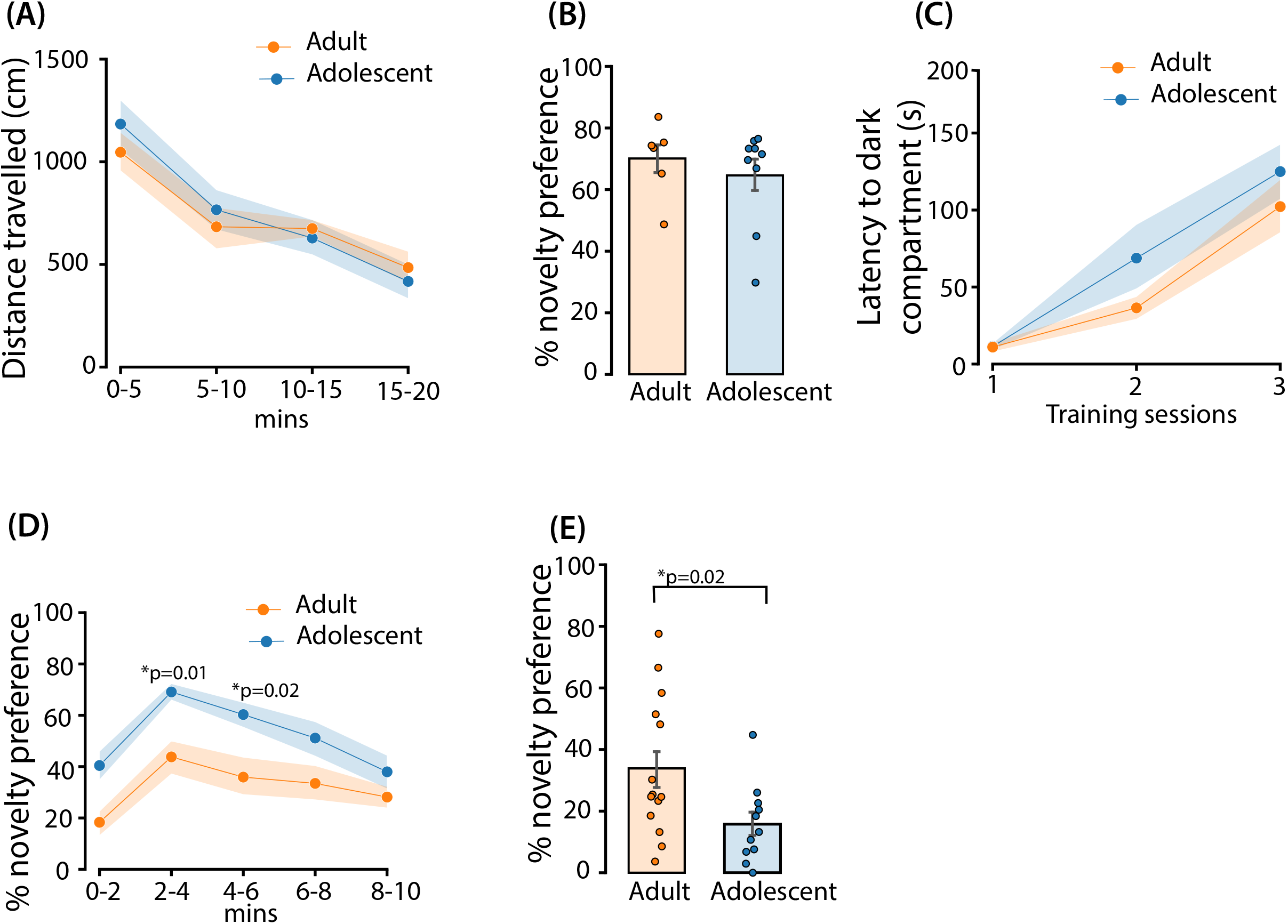

We then assessed adaptive behavioral coping strategies, assuming a possible measure of executive control and reward sensitivity ^36^, using operant conditioning paradigms. We first observed a similar capacity to self-control in absence of reward availability after a long training for saccharine self-administration. Specifically, although adult rats exhibited enhanced responding on the active lever for reward when saccharine was available, most likely reflecting their higher ingestion capacity, both groups significantly decreased their reward seeking behavior during periods of signaled unavailability (Figure 2A). Ultimately, response inhibition (reflecting the drastic decrease in lever presses when reward is no longer available) was similar between groups emphasizing that juvenile and adult rats exhibited comparable levels of response inhibition in the signaled absence of reward (Figure 2B). Further challenging their active behavioral responses, we found a similar level of motivation on an effort demanding reward-seeking task as indicated by the total number of lever presses (Figure 2C) and the total number of rewards (Figure 2D) earned during a progressive ratio schedule of reinforcement.

**Figure 2.**
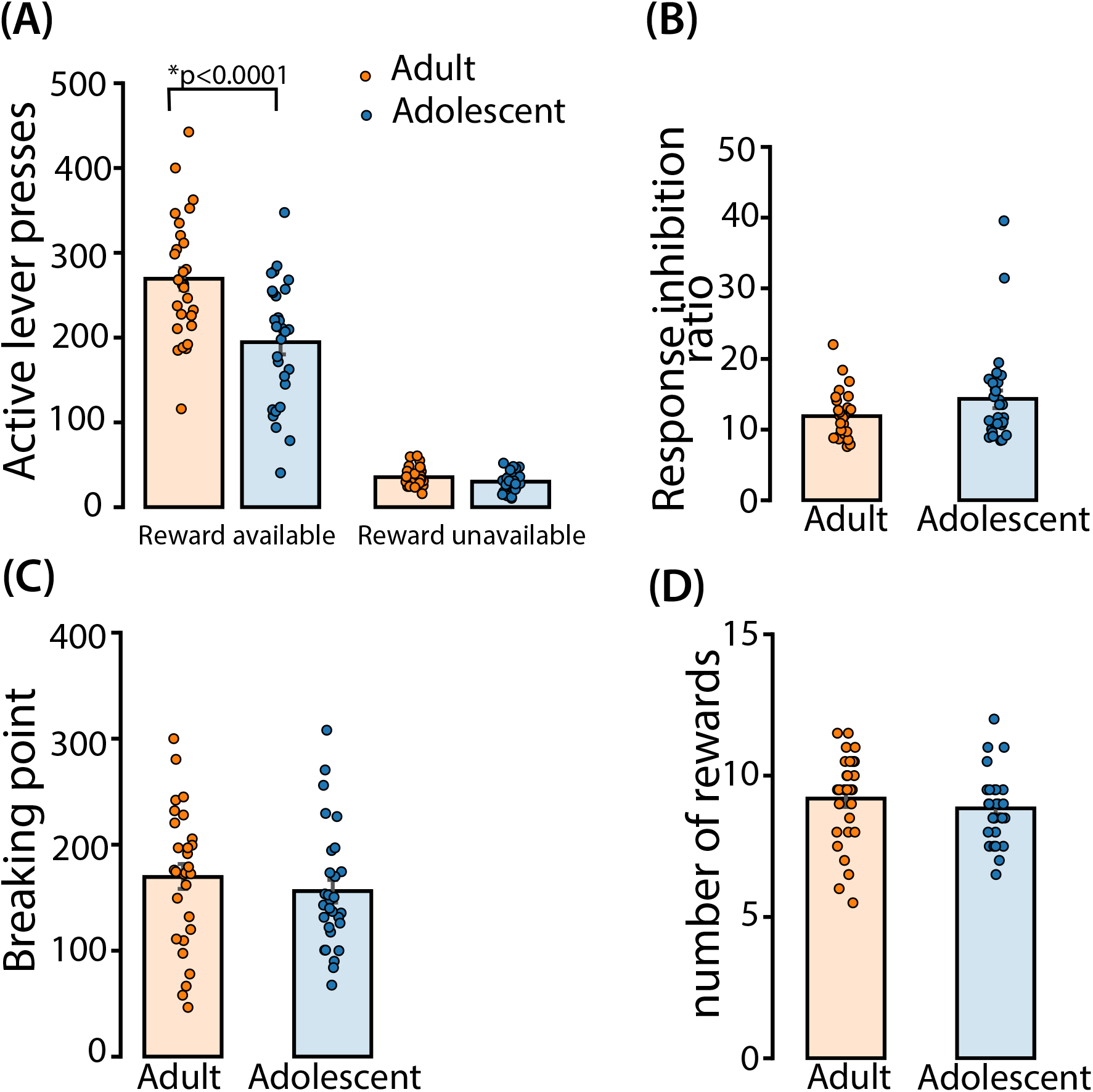

Overall, these observations indicate that juvenile and adult rats manifest rather comparable general coping strategies in absence of conflictual decision making, with larger fear responses in juveniles exposed to physical threat.

### Juvenile rats display persistent aberrant reward taking behaviors in conflictual situations reflecting enhanced impulsive- and compulsive-like behaviors

Converging evidence suggests that the maturation of brain regions supporting response inhibition may underlie the decline in risk taking behavior observed from adolescence through adulthood ^37^. Here we first tested juvenile and adult rats in a 5-choice serial reaction time task to assess their adaptive strategy (and response inhibition capacity) when sucrose food pellet delivery depends on behavioral performance contingent on stringent rules where withholding the behavioral response is mandatory and breaking the rule is counterproductive. Rats were trained and ultimately tested while food restricted and tested again while fed ad libitum. After extensive training sessions, juvenile and adult rats exhibited comparable performance reflecting similar response accuracy (Figure 3A) and limited omissions (Figure 3B), whether they were satiated or not.

**Figure 3.**
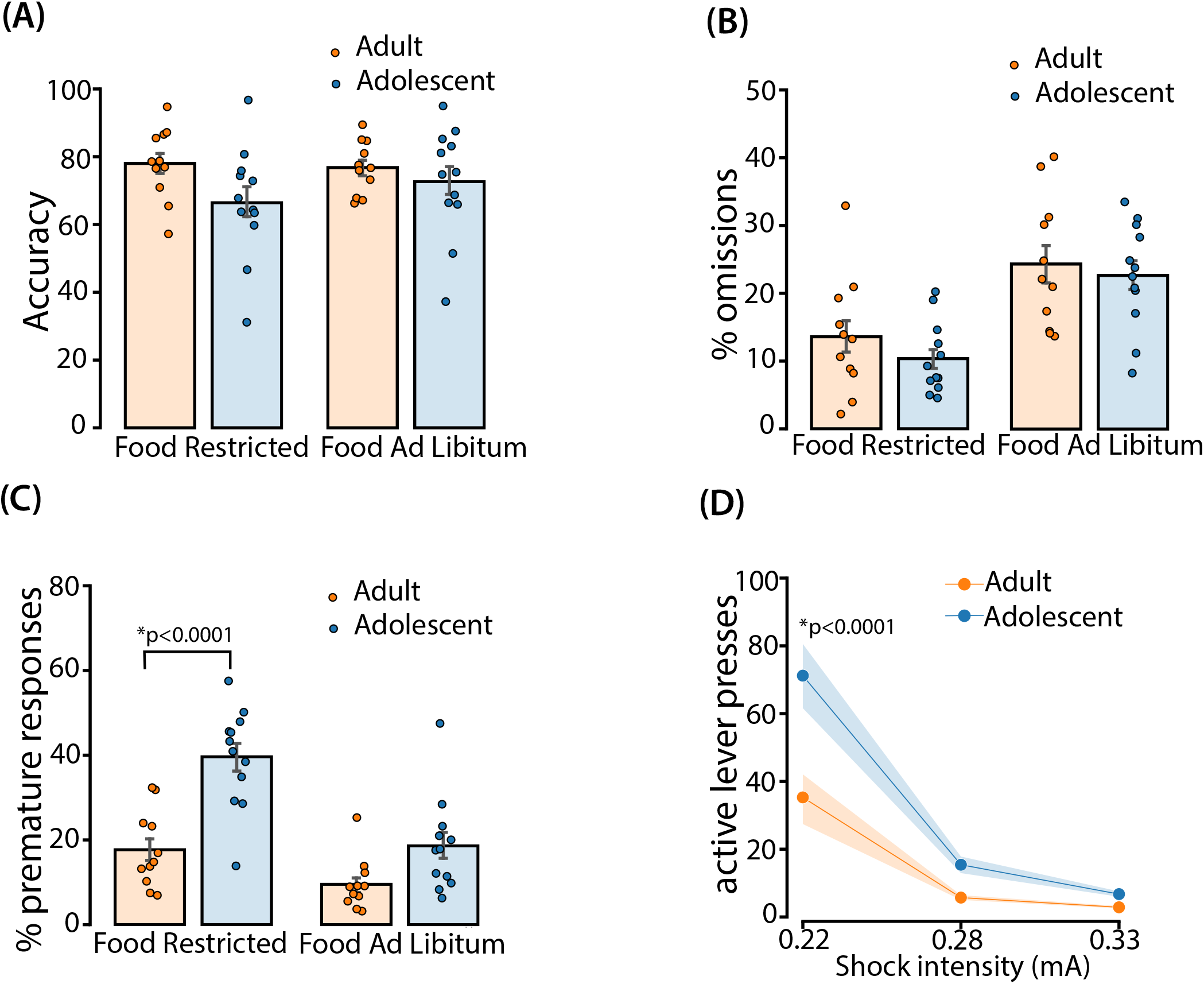

However, adolescent rats made more premature responses than adults (Figure 3C). After adjustment for multiple comparisons, juvenile rats expressed higher premature responses compared to adults upon food restriction, but significant differences between groups were not observed when rats were fed ad libitum. These observations suggest that in a conflictual situation (withholding a behavioral response while hunger promotes food seeking), juvenile rats encounter higher difficulty to adapt and manifest a counterproductive impulsive-like response.

Following these observations, we further challenged juvenile and adult rats using a conflict paradigm in which reward delivery was followed by punishment. Here, rats fed ad libitum had to adapt their lever pressing behavior when 0.2% saccharine solution delivery was followed by an electrical foot shock that increased in intensity over 3 consecutive sessions. As expected, lever presses decreased with increased shock intensity from 0.22 to 0.33 mA in all rats, but to a significantly smaller extent in juveniles. Strikingly, the adolescents persisted in lever pressing despite 0.22mA mild electrical foot shock, suggesting a compulsive-like reward seeking behavior (Figure 3D). Importantly, this effect could not be attributed to differences in nociceptive thresholds between adolescent and adult rats (Supplementary figure 1).

#### Lower functional recruitment of the anterior insula in juvenile rats compared to adults, not in the prelimbic cortex

To investigate potential mechanisms of differences in compulsive responding between adult and adolescent rats we examined the expression of mRNA for the transcription factor zif268 in the prelimbic cortex (PLC) in line with the reduced top-down executive control theory ^10, 24^ (Figure 4A). Although we did not find any difference in zif268 mRNA expression in PLC, we did observe a significant decrease in AIC (Figure 5A).

**Figure 4.**
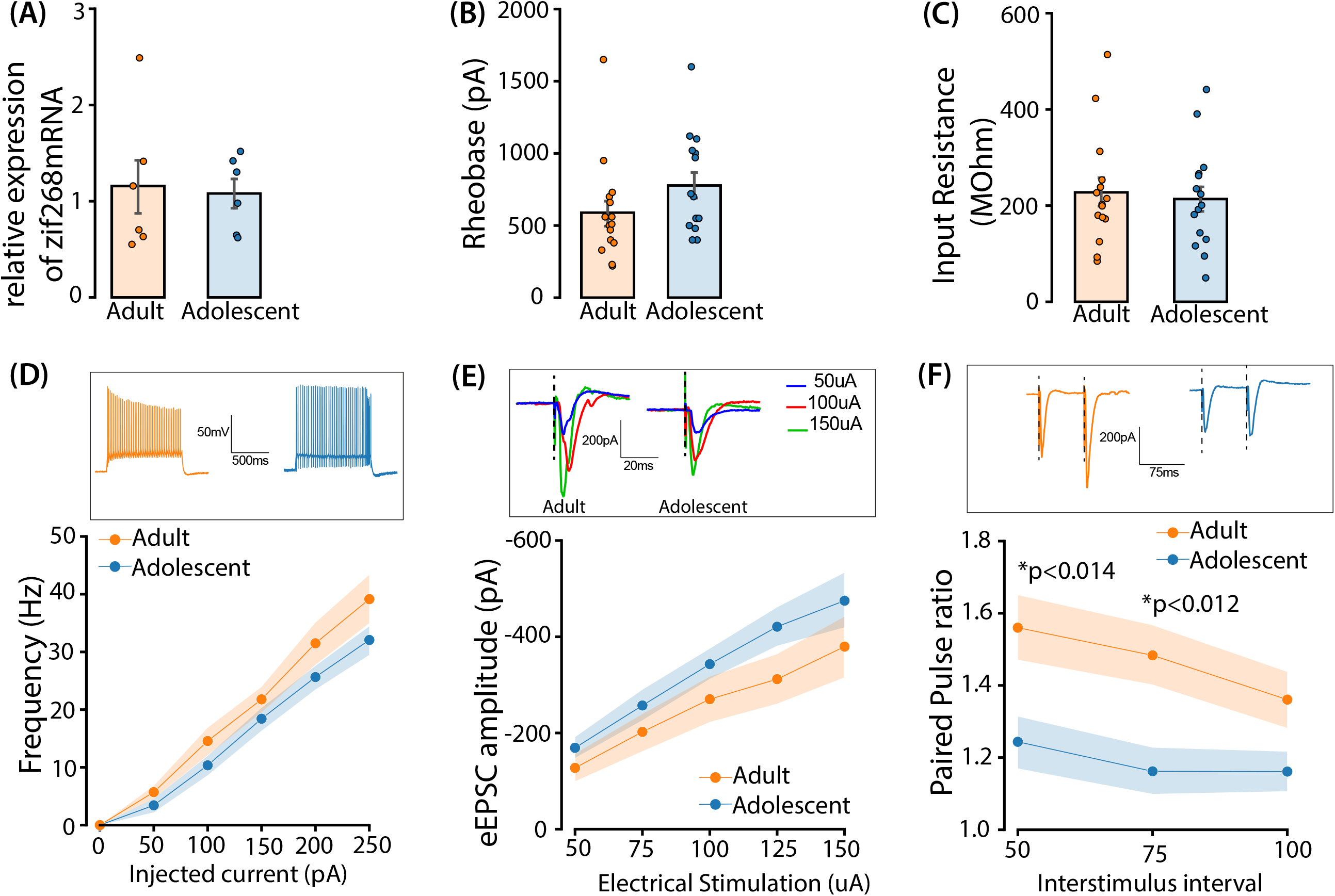

**Figure 5.**
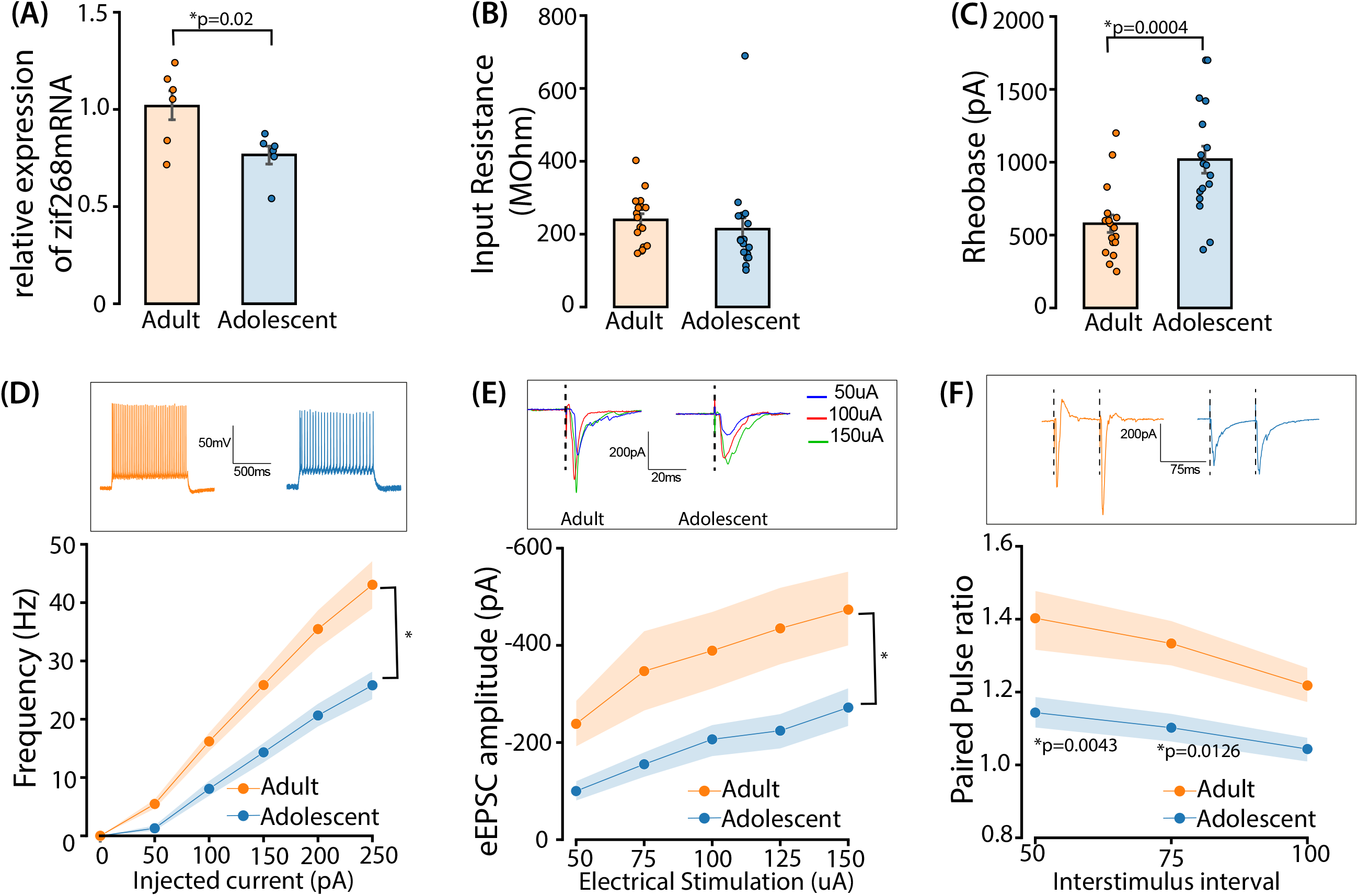

Our next experiments examined whether differences in the functional properties of layer 5 (L5) PLC or AIC pyramidal neurons exist between adolescents and adults. The intrinsic excitability and strength of glutamatergic inputs impinging on L5 pyramidal neurons was assessed using *in vitro* whole-cell recordings, and synaptic currents were evoked using local electrical stimulation. Pyramidal neurons were identified using electrophysiological criteria, such as the lack of spontaneous action potentials after gaining whole-cell access, and the presence of action potential accommodation upon current injection.

We found that rheobase (Figure 4B) and input resistance (Figure 4C) of L5 PLC pyramidal neurons was not significantly different between adult and adolescent rats. Moreover, these neurons also exhibited similar action potential discharge rates with a series of current injections (Figure 4D), and the mean amplitudes of electrically evoked EPSCs (eEPSCs) did not differ between adolescent and adult rats (Figure 4E). Interestingly however, a smaller paired pulse ratio at inter-stimulus intervals of 50ms and 75ms was observed in the adolescent group indicating a possible decrease in the probability of glutamate release from the presynaptic projections on to the PLC L5 pyramidal neurons (Figure 4F).

Like the PLC, AIC L5 pyramidal neuron input resistance did not differ between adult and adolescent rats (Figure 5B). However, unlike the PLC, the rheobase of the AIC L5 pyramidal neurons was significantly higher in the adolescent group as compared to adults (Figure 5C). This decrease in threshold excitability was accompanied by the diminished excitability of these neurons when the number of action potentials initiated by current injection was found to differ between adolescents and adults (Figure 5D). We also found that the relationship between stimulus intensity and eEPSC amplitude was significantly smaller in adolescent AIC L5 pyramidal neurons, compared to adults (Figure 5E). Moreover, we also observed smaller paired pulse ratios in L5 pyramidal neurons from adolescents at inter-stimulus interval of 50ms and 75ms, suggesting that the probability of glutamate release from the axonal projections onto AIC L5 pyramidal neurons was smaller in the juvenile group (Figure 5F).

### Chemogenetic activation of the AIC attenuates compulsive-like behavior in juvenile rats

As our electrophysiological studies indicated that L5 AIC neurons were hypoexcitable and received less excitatory input in adolescent rats, we next examined whether bilateral chemogenetic activation of the AIC using the hM3D construct would decrease foot shock resistant reward taking in adolescent rats (Figure 6A and 6B). We found that the DREADD agonist clozapine had no effect on basal performance either in sham controls or in rats injected with AAV-hM3D (on PND 45-46, Supplementary figure 2). However, clozapine injection resulted in a significant reduction of saccharine taking during punished responding in the adolescent rats that had received AIC injections of the excitatory DREADD viral construct (Figure 6 C and D).

**Figure 6.**
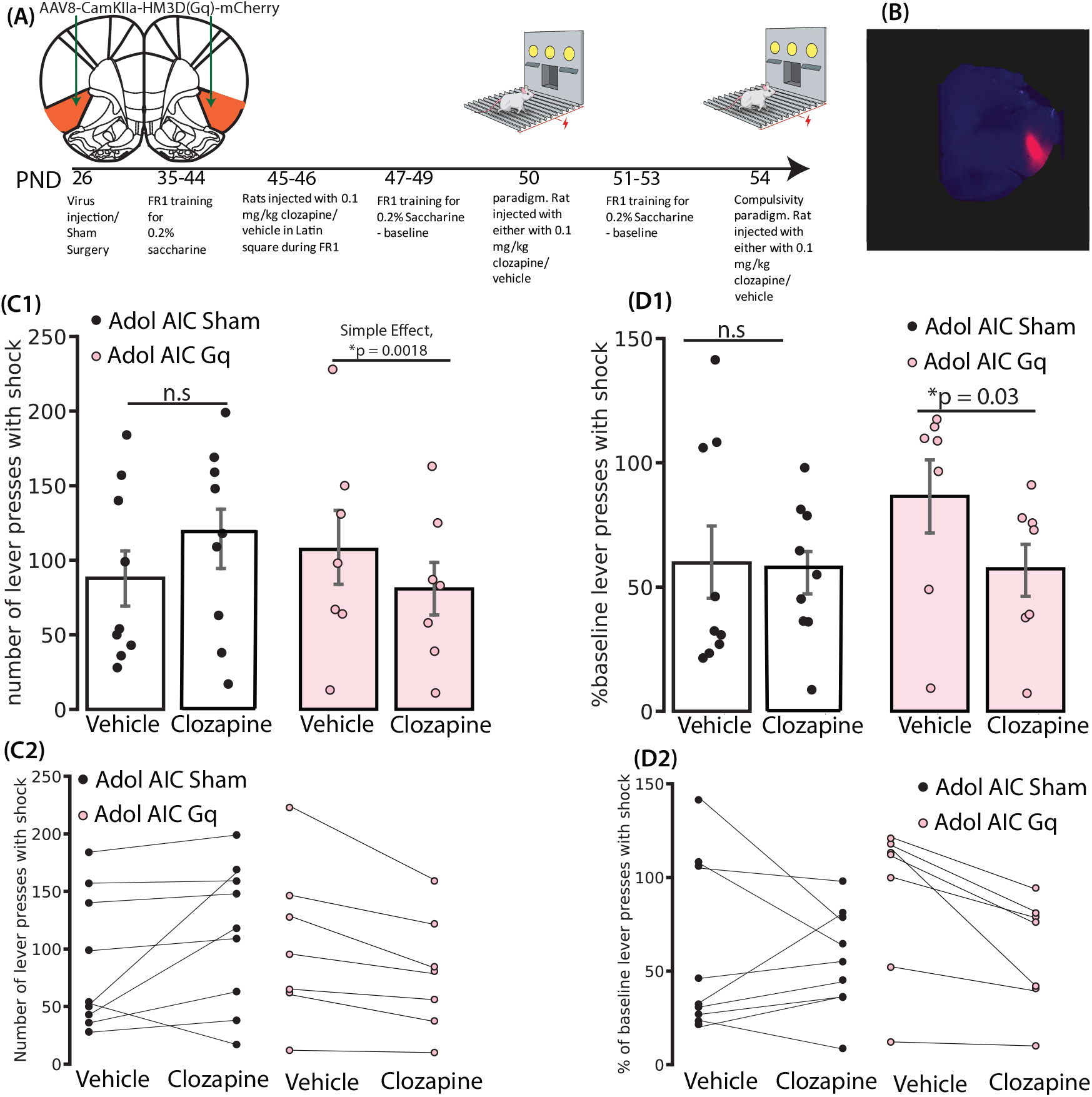

Altogether, these results indicate that chemogenetic activation of the AIC in juvenile rats does not disrupt basal performance in an operant conditioning task in absence of foot shocks, but significantly reduces persistent lever pressing behavior during conflictual decision making, precisely when a reward is paired with a punishment.

## Discussion

In the present series of experiments, we found that whereas juvenile rats exhibited most of the general coping strategies present in adults, they also show a much higher degree of inflexible behavior when making decisions under conflict. We also observed a decrease in Mrna expression of zif268 in the AIC of the adolescent rats and found using *in vitro* electrophysiology decreased intrinsic excitability of L5 pyramidal neurons, as well as a weaker glutamatergic synaptic input to these cells. To determine whether this AIC functional immaturity could underlie increased compulsive behavior in adolescent rats we reversed the hypoexcitable state of the AIC by chemogenetic activation, and this significantly attenuated this behavior.

According to neural imbalance models of behavioral control, top-down inhibition of subcortical brain structures by cortical regions is weak during adolescence, promoting impulsivity, sensation seeking, and reckless behavior ^4, 6^. However, during maturation it may be advantageous to facilitate novel learning strategies through increased risk and novelty-seeking behavior ^38^.

Here, we found that juvenile and adult rats largely exhibited similar open field exploration and novelty place preference behavior (Figure 1A and 1B) ^38, 39, 40, 41, 42^. They also displayed similar drive for novel environment and comparable ability to control for saccharine seeking in its absence (Figure 2A and 2B), as well as comparable learning skills in the motor impulsivity paradigm (Figure 3A and 3B). However, juvenile rats did exhibit greater premature responses in the 5CSRTT during food restriction. This observation reveals their inability to self-regulate under a conflicted situation of increased drive to gain food, as already reported in the literature ^38, 43, 44^. It also confirms that the ability to regulate reward- and sensation-seeking behaviors may not depend solely on impulse control but may rely on subsequent sensory/cognitive processes. Supporting this interpretation, adolescent rodents exhibit better performance when seeking rewards in a cue-guided reversal procedure (with location of reward indicated by odors ^45^). They also perform better in tasks revealing cognitive flexibility through different reward contingencies ^46^. This enhanced flexibility in adolescence, considered as a prefrontal cortex-dependent function ^47, 48^, is rather counterintuitive to the idea that the PFC is considered functionally immature at this developmental stage and calls for reassessment of current interpretations ^3, 10, 12, 13^.

Our key observation is that adolescent rats exhibited higher persistence for saccharine reward despite negative consequences (Figure 3D). Impulsivity is considered an endophenotype for higher vulnerability to compulsive behavior in many clinical and preclinical studies ^30, 49, 50, 51, 52, 53, 54^, and is thought to reflect loss of control over reward-seeking in rodents ^55, 56^ and enhanced risk-taking in humans ^57, 58^. In the current study, we controlled for potential differences in the appetitive properties of saccharine (Figure 3C and 3D) and pain thresholds (Supplementary Figure 1) between adolescent and adult rats. We also controlled for emotional reactivity to pain or risky situations in juveniles and report similar passive avoidance, increased freezing behavior and increased time in the closed arm of the elevated plus maze (Figure 1C, 1D, 1E), mitigating the idea that differences in fear processing could explain higher risk taking in adolescent rats. We then revealed similar mRNA expression of the early gene marker zif268 in the PLC (Figure 4A), and confirmed similar intrinsic excitability of L5 PLC pyramidal neurons in all rats (Figure 4B, 4C, 4D, 4E). Considering that both the rodent medial PLC and primate dorsolateral PFC support executive functions ^59^, our observation challenges the theory of decreased top-down control of the immature prefrontal cortex as a possible explanation for adolescent compulsivity. In contrast to PLC, we found decreased expression of zif268 mRNA in AIC, and this was accompanied by a lower excitability of AIC L5 pyramidal neurons and weaker synaptic glutamate inputs to these neurons in adolescent rats (Figure 5B, 5C, 5D, 5E, 5F). This indicates that not only do these neurons receive weaker glutamatergic projections, but that they are also less-responsive to these inputs, reflecting reduced ability to integrate information. These findings confirm that increased sensitivity to punishment from adolescence to adulthood may be subserved by greater neural recruitment of the insula ^31^.

The relationship between the neuronal hypoexcitability of the AIC in increased compulsivity in adolescents that we describe here are correlational. Therefore, we used chemogenetics to increase the activity of the AIC in juvenile rats during punished reward seeking to determine whether the reversal of hypoactivity could affect behavior. We found that chemogenetic stimulation of the AIC in adolescent rats significantly attenuated persistent punished reward taking behavior (Figure 6) without affecting motor performance (supplementary figure 2). This suggests that the diminished excitability of AIC L5 pyramidal neurons and the smaller synaptic glutamate drive of these cells likely contribute to differences in AIC-dependent behavior between adolescent and adult rats.

The decreased synaptic glutamatergic strength together with the hypoexcitability of AIC pyramidal neurons could represent a mechanistic explanation for the lower ability of adolescent rats to integrate sensory, emotional and cognitive information ^28, 29^, and this might explain why adolescent rats are less able to process converging interoceptive signals, particularly aversive ones, in risky decision making tasks, as has already been suggested in humans ^60, 61, 62^. The reduced functional activity observed in AIC contrasts with similar activity in the PLC of adult and adolescent rats may reflect the asynchrony of maturation of these cortical structures and suggests that the connections between these two structures are not fully established in adolescent rodents.

A recent hypothesis suggests that the insula computes interoceptive predictions by estimating both current and future physiological needs (i.e., a change in an expected interoceptive state) to guide brain and behavior toward a homeostatic set point ^63^. In this model, rather than the simple integration of sensory inputs, interoceptive perceptions can be viewed of as Bayesian estimation of past experience (inference) about the sensory consequences of homeostatic budgeting that are implemented as upcoming visceromotor signals ^63, 64^. These inferences are also constrained by error signals that result from the failure of previous predictions to accurately account for incoming interoceptive sensations ^65^. Ultimately, not only does past viscerosensory experience influence the present experience, but the present one project forward to influence what will be perceived in the future, meaning that interoceptive perception is largely a construction of beliefs constrained by the actual state of the body ^66^. This fundamental assumption supports the idea that the insula makes interoceptive predictions and prediction errors that could be considered risk prediction and risk prediction errors. Supporting this interpretation, a late-onset anticipatory risk prediction signal followed by a fast-onset prediction error signal at the time of the outcome have been reported during a gambling task in humans ^67^. Considering this bi-modal response within the AIC for prediction learning, the long-lasting effect of the chemogenetic activation we report here might have partially biased the expected substantial lowering of compulsive lever pressing in adolescent rats (Figure 6). Another comment arising from this assumption regarding the need to minimize the difference between the brain’s prediction and incoming sensation is the issue of how and when specific bodily signals are consciously represented, in adolescent and adult brains. The neuronal hypoexcitability reported here in the AIC of juvenile rats opens an interesting debate about how the risk is integrated (awareness) but not learned optimally (consciousness) ^27^, perhaps explaining the persistent lever pressing despite punishment observed in juveniles. Considering this transient functional anosognosia (that we would define as an operating self-awareness but not yet a full consciousness), we would like to suggest that the maturation of the AIC contributes to the construction of brain positive (and negative) alliesthesia ^68^ in the adolescent brain. In line with the role attributed to the AIC in the active inference framework, perception and action are tightly coupled, according to brain’s estimations following the integration of both extero and interoceptive cues. Interoceptive perceptions therefore derive from the brain’s best prediction to infer the causes of the sensations it receives, constrained by incoming sensory inputs. Either the adolescent brain accepts the punishment as a prerequisite for accessing the reward when the adult considers it is not worth the behavioral output, or the anticipated lack of reward is integrated as an even more painful punishment as compared to the foot shock itself, and the persistent lever pressing behavior is motivated by the expected positive affective state associated with reward consumption. Our observations only partially unveil how the delayed maturation of the insula may result in suboptimal integration of sensory and cognitive information in adolescents, possibly promoting inflexible reward-related behaviors. But understanding how interoceptive inference contributes to shaping the emotional brain, notably during adolescent development, is of the highest relevance, in particular for preventing the emergence of psychiatric conditions.

## Author contributions

Conceptualization, KSJ, BB; Methodology, KSJ, APB, AFH, ID, CRL, BB; Behavior analyses, KSJ, APB, LA, GK, ID, BB; Pain sensitivity, GK, ID; Ex vivo patch clamp electrophysiology, KSJ, AFH, CRL; In Vivo Chemogenetic studies, KSJ, LA, BB; First draft writing, KSJ, BB; Review and comments on draft, CRL, APB, ID; Funding acquisition, CRL, BB; Project supervision and administration, BB.

## Acknowledgments

The authors would like to thank Nathalie Habegger, Jennifer Silva and Leticia Pedrido for technical assistance, Prof Olivier Halfon for constructive comments, and Cédric Leser for technical assistance in the cartoon drawing on figure 6. This work was supported by the Department of Psychiatry, Lausanne University Hospital and a Swiss National Science Foundation grant (310030_185192 to BB), and NIH Intramural Research Grant 1ZIADA000457 to CRL from the US Department of Health and Human Services. KSJ is recipient of a Swiss Government Excellence Scholarship, a Doc mobility fellowship from the Swiss National Science Foundation, and the 2020 best thesis award from the Société Académique Vaudoise.

## Data availability

The data that support the findings of this study are available from the corresponding author upon reasonable request.

## Material and methods

### Animals

All experiments conducted in Switzerland used male Wistar rats (130 adolescents and 102 adults) from our breeding colony (breeders ordered from Charles River, France). Adolescent rats (PND 21) weighed about 70-100g and adult rats (PND 70) weighed 250-300 g at the beginning of experiments. Rats were grouped 3 per cage (560×330×270 mm). They were kept under a 12/12 h light/dark reversed cycle (light off at 8:30AM), in a room at 22 ±2 °C and 40-60% humidity. Each behavioral test has been conducted during the dark phase of the cycle. Except in some specific conditions, animals had ad libitum access to water and food (standard chow, 3436, Kliba, Provimini, Switzerland). All procedures described were conducted in accordance with the Swiss National Institutional Guidelines on Animal Experimentation and approved by the Swiss Cantonal Veterinary Office Committee for Animal Experimentation (authorization 1999.3, 1999.4 and 3047.1b to B.B).

For the electrophysiology experiments 6 adolescent and 6 adult male Wistar rats were ordered from Charles River, USA. Rats were grouped 3 per cage (560×330×270 mm). They were kept under a 12/12 h light/dark reversed cycle (light off at 8:30AM), in a room at 22 ±2 °C and 40-60% humidity. The electrophysiology experiments were conducted at P50 for adolescent rats and P90 for adult rats.

### Five-choice serial reaction time task (5CSRTT)

We used six operant chambers (305 × 241 × 292 mm, Med Associates Inc., St. Albans, VT, USA) individually enclosed in a wooden cubicle, equipped with an exhaust fan for ventilation as well as served as a white noise speaker for sound attenuation. On the left side of the chamber, five cavities (25×25 mm) for the nosepokes were present horizontally and separated by 25mm from each other. Opposite these five apertures, on the right side of the chamber, was the food receptacle, located 20mm above the grid. Lights for the nosepokes, food receptacle and chamber ceiling were present. Sucrose pellet (Dustless precision pellet 45mg, rodent purified diet, Bioserv, Frenchtown, NJ) were used during the whole experiment as a reward. The 5CSRTT was carried out following five training steps explained in supplementary information.

### Operant conditioning

Experiments were conducted in six operant chambers (305 × 241 × 210 mm, Med Associates Inc., St. Albans, VT, USA) placed in sound-attenuated cubicles equipped with ventilation fans. Each operant panel had two automatically retractable levers, located 70 mm above the grid and 35 mm equidistant from the midline. A white light diode was mounted 30 mm above each lever. Between both levers, a food tray was linked to a pellet delivery system. The food tray further included a cavity that allowed liquid solution delivery through a dipper system. The operant training was carried out following several training steps explained in supplementary information.

### Early gene marker zif268

Six adolescent and six adult rats were trained on the persistence in reward seeking despite 0.22mA electrical foot shock punishment and were sacrificed 15 minutes after the end of the 30-minute session. Following rapid decapitation and extraction, the brains were sliced at 2mm thickness using the brain matrix. The slices were immediately frozen on glass slides using dry ice. The Prelimbic cortex, AIC and DLS were punched using 1 mm punches. RNA was extracted with a ReliaPrep RNA Tissue Miniprep System (Promega, USA) and converted into cDNA by reverse transcription reaction using TaqMan Reverse Transcriptase Reagents (Applied Biosystem, Foster City, CA, USA). Real-time PCR amplification was performed with an ABIPRISM 7500 cycler and SYBER green PCR Master Mix (Applied Biosystem, Foster City, CA, USA) using specific sets of primers (Microsynth AG, 9436 Balgach, Switzerland). Forward and reverse primers for the zif268 gene are the following: β-actin = forward: 5’-TCTGAATAACGAGAAGGCCGTGGT-3’ and, reverse: 5’-ACAAGGCCACTGACTAGGCTGAAA-3’. All samples were analyzed in triplicates. Relative gene expression was measured with the comparative ΔΔCt method24 and normalized with β-actin transcript levels.

### Slice electrophysiology

#### Brain Slice Preparation

Animals were anesthetized with isoflurane and decapitated using a guillotine. Brains were extracted and transferred to an ice-cold cutting solution containing (in mM): N-methyl-D-glucamine (NMDG), 93; KCl, 2.5; NaH2PO4, 1.2; NaHCO3, 30; HEPES, 20; Glucose, 25; Ascorbic acid, 5; Sodium pyruvate, 3; MgCl2, 10; CaCl2, 0.5. The brain tissue was cut perpendicular to its longitudinal axis using a razor blade, and then the cut surface glued to the stage of the vibrating tissue slicer (Leica VT1200S, Leica Biosystems). Coronal sections (250 μm) were collected from each brain. The brain slices were then transferred to an oxygenated (95% O2/5% CO2) holding chamber filled with HEPES containing artificial cerebrospinal fluid (aCSF) (in mM: NaCl, 109; KCl, 4.5; NaH2PO4, 1.2; NaHCO3, 35; HEPES, 20; Glucose, 11; Ascorbic acid, 0.4; MgCl2, 1; CaCl2, 2.5) at room temperature (22°C) for at least 30 min.

#### In Vitro Electrophysiology

Hemi-sectioned slices were transferred to a recording chamber (RC-26; Warner Instruments) continuously perfused (2 mL/min) with oxygenated aCSF (in mM: NaCl, 126; KCl, 4.5; NaH2PO4, 1.2; NaHCO3, 26; Glucose, 11; MgCl2, 2; CaCl2, 2.5) using a peristaltic pump (Cole-Parmer). The temperature was maintained at 30–32°C using an inline solution heater connected to a temperature controller (TC-344C/SH-27B, Warner Instruments). Appropriate neurons were identified using an upright videomicroscope (BX51WI, Olympus) equipped with differential interference contrast imaging, and 900 nm infrared illumination. Recording electrodes were fabricated using borosilicate glass (Sutter Instruments, 1.5 mm O.D. × 0.86 mm i.d.) using a horizontal puller (P-97; Sutter Instruments) and filled with a potassium-based internal solution (in mM: K-gluconate, 140; KCl, 5; HEPES, 10; EGTA, 0.2; MgCl2, 2; Mg-ATP, 4; Na2-GTP, 0.3; Na2-phosphocreatine, 10) neutralized to a pH of 7.2 using potassium hydroxide. Electrode resistances were 3–5 MΩ. Whole-cell patch clamp recordings were performed using an Axopatch 200B amplifier (Molecular Devices). Unless otherwise noted, cells were voltage-clamped at −60 mV (with a calculated junction potential of −14 mV). Stimulation protocols and recordings were performed using WinLTP software (WinLTP Ltd, Bristol, UK) and an A/D board (National Instruments, PCI-6251) housed in a personal computer. Hyperpolarizing voltage steps (−10 mV) were delivered via the recording electrode every 30s to monitor whole-cell access and series resistance. Cells with an access change of more than 20% were excluded from analysis.

#### Input resistance

Hyperpolarizing voltage steps (−10 mV, 200 msec) were delivered via the recording electrode every 30 s to monitor the steady state input resistance, obtained over the last 20ms of the step. The input resistance was calculated as the average of the last 10 sweeps in voltage clamp.

#### Rheobase

Rheobase (the lowest current injection to elicit a single action potential) was evaluated in current clamp. The depolarizing current steps were applied (0–2000pA, 50pA increments, 5ms pulse duration, repeated 3 times) from a membrane potential of -60mV.

#### Excitability

To measure excitability, depolarizing current steps were applied (0–250pA), 50pA increments, 1s pulse duration, repeated 3 times from a membrane potential of -60mV. The mean number of action potentials (APs) per second during each depolarization step was plotted against the amount of current injected.

#### Electrically elicited EPSCs

Excitatory synaptic currents were evoked using bipolar tungsten stimulating electrodes with a tip separation of 300-400 μm. Stimulating electrode tips were placed within 1 mm rostral of the recording patched neuron. Whole cell access was monitored using voltage (or current) step pulses (−10–20 mV, 200 ms) delivered after each stimulus using the Master-8 pulse generator. Stimulation (0.1-ms pulse duration) was delivered at 30-s intervals using an optically isolated constant current unit (AMPI) and the timer. Input–output (I-O) relationships between elicited EPSC amplitude and electrical stimulation intensity (50, 75, 100, 125, 150 uA) were constructed. Paired-pulse stimulation was performed by delivering the same stimulus at 50 ms, 75ms and 100 ms (EPSC) inter-pulse intervals.

### Chemogenetics

Adolescent rats at postnatal day (PND) 26 were used for viral injection in the anterior insular cortex. Rats were anesthetized with isoflurane using a precision vaporizer connected to a nose cone apparatus and injected (0.8 μl over 15 min) with AAV8 excitatory DREADD (pAAV-CaMKIIa-hM3D(Gq)-mCherry) (4.6×1012 virus molecules/ml; University of North Carolina vector core) or into the anterior insular cortex (AP: +2.4, ML: +-4, DV: -5.2), using a 5-μl Hamilton syringe, an UltraMicroPump, and SYS-Mico4 controller (WPI, Sarasota, FL). Incisions were closed with absorbable sutures, and body temperature was maintained with a heating pad until anesthesia recovery. Rats were given 1 week to recover and then trained on the Fixed Ratio 1 schedule of reinforcement training procedure as mentioned before for a period of 15 days followed by testing their compulsivity for sweetened taste in presence of mild foot shock (0.22mA). Clozapine (0.1mg/kg ip) or Vehicle (equivalent volume) were injected intraperitoneally 30 min before the compulsivity test and the rats were tested in a Latin Square design so that each rat could serve as its own control. Post-mortem verification revealed that 1 rat received the active virus in a brain site off target and hence was eliminated from further analysis.

### Statistical Analyses

Data shown are represented as group mean ± SEM. For single group comparison, Student t-tests were performed for normal distributions, or non-parametric Mann-Whitney U-tests for non-normal distributions. ANOVAs were conducted for repeated measurements, but non-linear mixed effect models were also used to deal with missing values or non-normal data distribution. Appropriate Post-hoc Sidak’s tests were further used for identifying group level differences. All statistical analyses were conducted using the statistical software GraphPad Prism (Version 6.07). Significance of results was accepted at p < 0.05.

## Supplementary information

### Locomotor habituation and novelty preference

The arena was divided in two chambers (45×46×30cm) linked by a central compartment (39×15×30cm). Both chambers differed by the ground texture and wall patterns. Nine adolescent and seven adult rats were used for this experiment. Rats were first placed in the central compartment for 5 min. Then, they were put in one of the two other compartments for 20 min, in a counterbalanced fashion. During this familiarization phase, the door stayed closed and locomotor activity was monitored. Immediately after, rats were put in the center compartment, and both doors were removed. Locomotor activity on the first 20 min was used as a measure of locomotor habituation, and time spent in each compartment during the second phase allowed to calculate preference ratio for novelty.

### Control for the emotional fear assessment

#### Contextual fear conditioning

We used an operant chamber (318×241×210 mm) bordered by operant panels all over, with a cue light at the top of one side and a tone device on the other. Nine juvenile and ten adults were conditioned for two days and tested on the third day. Conditioning sessions lasted 20 min. During the first 10 min, no shocks were delivered. Then, 8 shocks (0.55 mA) of 1 sec were delivered, with variables inter-shock intervals to promote unpredictability. For the testing day, no shocks were delivered, and the freezing time was recorded using the video tracking system and manually scored for 10 min.

#### Passive avoidance

The apparatus consisted in a cage (440×171×254 mm), divided in the middle by an automated door (Med associates, Inc, St. Albans, Vermont, USA). Each side was completely identical, with a light device at the extremity. Four infrared beams were located in each chamber, 1 cm above the grid floor, allowing detection of the rat. Nine juvenile and nine adults were used. The protocol consisted in a multi-trial acquisition of the passive avoidance response, followed by a 24-h retention test. During the acquisition phase, a trial started when the rat was put in the illuminated chamber. Twenty seconds after, the door opened for 160 sec. If the rat stayed in the illuminated compartment, the door closed, the rat was brought out from the cage and the acquisition phase was finished. If the rat moved into the dark compartment, the door closed, and the rat received a 5-sec 0.22 mA foot shock. Then, he was moved off the cage, and was returned into it 60 sec after. This training was repeated up to a maximum of 3 times. Rats were tested again 24 hours later for a single trial retention test in the same conditions. Entry latency to the dark compartment was used as an index of emotional reactivity.

#### Elevated plus-maze

The arena was positioned 50 cm above the floor and divided in four arms: two “closed arms” enclosed by plastic wall (500×100×425 mm), and two “open arms” without walls. In the center, a small open arena (100×100 mm) allowed access to each arm. Luminosity was fixed at 20 lux in the open arms, and 5 lux in the closed arms. Rats were placed at the center of the arena, and their exploratory activity was recorded for 5 min. Percentage of time spent in the open arms was used as the index of anxiety, and locomotor activity was further assessed.

### Five-choice serial reaction time task training sessions

*For Training 1*, rats were trained to perform a nosepoke in the food receptacle to receive one sucrose pellet until they reached 50 pellets in 30 mins.

*For Training* 2, after the rat made a nosepoke in the food receptacle, a light stimulus appeared randomly in one of the five receptacles until the rat explored the illuminated receptacle. The light turned off then and the rat received a pellet in the food receptacle. Random nosepoking in unlit receptacles had no consequence. Rats were trained until they earned 40 pellets in a 30-minutes training.

*Training 3* was similar to Training 2, with a delay of 5 seconds introduced after the beginning of the trial and before one of the receptacles was illuminated. Rewarded correct (in the illuminated receptacle) and unrewarded incorrect (in any of the other four unlit receptacles) responses were recorded. A new trial began following a nosepoke in the food receptacle, a 5 second waiting period and a random nosepoke that was lit. Rats had to learn this step until they reached 50 correct responses during the 30minute training.

*For Training 4*, each trial following a nosepoke in the food receptacle was signified by a sound tone during 5 seconds. After the tone, one of the receptacles was illuminated for a period of 2 seconds. A response during the tone and before a receptacle being illuminated was measured as a premature response but there were no programmatic consequences. Correct and incorrect responses were recorded as well as omission responses, (no response during the allowed 5 seconds after the nose poke was lit). To advance to the final test session (during which premature responses were unrewarded and signaled by home light illumination for 5 seconds before next trial, see Figure A), the rats had to make at least 30 correct responses during the 30-minute session. Rats were tested while food restricted over 3 consecutive sessions and tested again over 3 consecutive sessions while fed ad libitum.

**Figure A:**
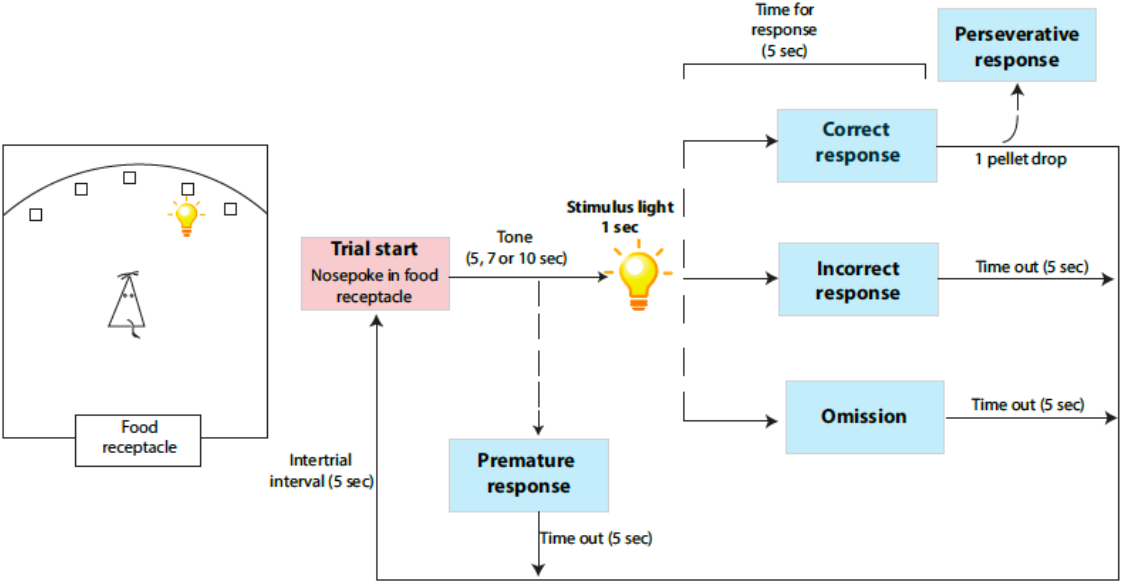
Pictorial representation of 5-Choice serial reaction time task test

### Operant conditioning

Rats were trained to self-administer 0.1mL of saccharine 0.2% liquid reward (Sigma-Aldrich Chemie GmbH, Buchs, Switzerland) on a fixed ratio 1, time out 4 sec (FR1 TO4) schedule of reinforcement during 30-min daily sessions. A single lever press on the active lever activated a liquid dipper equipped with a 0.1ml cup. The liquid reward remained available for 4 sec following lever-pressing before the dipper stepped down. Supplementary activity on the active lever during the time out period, as well as any lever presses on the inactive lever were recorded but had no further consequence. After stabilization of performance (±20% variation of the mean responses for three consecutive sessions), rats underwent 3 distinct test phases, with saccharin training sessions between each test to ensure that a test phase did not affect later saccharin consumption.

#### Control of reward-seeking

Twenty-nine adolescent and twenty-nine adult rats were used for these experiments. Here, 8 min of classical saccharin self-administration were followed a 4 min period of reward unavailability, signaled by the illumination of the house light and the shutdown of the ventilation fan. This cycle was repeated 3 times (for a total of 36 min), and total number of lever presses and rewards earned have been separately recorded and analyzed for each period.

#### Progressive ratio

The same twenty-nine adolescent and twenty-nine adult rats were used for evaluating their effort related motivation. The formula used to calculate increase of response requirements was: response ratio = 5 x e (injection number x0.2) – 5. The different ratios were therefore 1, 2, 4, 6, 9, 12, 15, 20, 25, 32, 40, 50, 62, 77, 95 and so on. The session ended either after 90 min, or after 30 min without any lever press. Lever presses rewards earned were recorded. The total number of active lever presses at which the rat stops pressing the lever is termed as the breaking point and is a measure of its motivation to gain the reward.

#### Persistence of reward-seeking

The same twenty-nine adolescent and twenty-eight adult rats were used for these experiments. One adult rat was deleted since the rat showed weight loss and reached the exclusion criteria. For this test, electric foot shocks were paired with reward delivery. After a lever press, cue light was on, and saccharin solution was available for 4 sec. A 0.5-sec foot shock occurred after reward availability. Three daily sessions were performed, with increased intensity of shocks: 0.22, 0.28 and 0.33 mA.

### Control for pain sensitivity thresholds

#### Pinch test

The pinch test was performed using a pair of large blunt forceps (15 cm long; flat contact area: 7 mm × 1.5 mm with smooth edges) attached to a 2 strain gauges connected with a modified electronic dynamometer (Bioseb). The forceps tips were placed around the rat paw or tail, and incremental force was applied till a withdrawal response. This was repeated a total of 3 times, and the mean force (in grams) which induced a withdrawal of that particular appendage was calculated.

#### Hot plate test

The rats were individually placed on the hot-plate surface which was set at defined temperatures (49°C, 52°C, and 55°C). The time until the response (latency in seconds) defined by a hind paw lick or jump was measured. The following cutoff times were used to avoid tissue damage (60 seconds for 49°C, 30 seconds for 52°C, and 20 seconds for 55°C)

#### Tail-flick test

A tail-flick analgesia meter (Columbus Instruments) was used to conduct this experiment. The rats were restrained gently using a conical plastic cloth. Two different light beam intensities [4 and 7 (AU)] were used and the latency of response (in seconds) was recorded.

## Legends for supplementary figures

**Supplementary figure 1:**
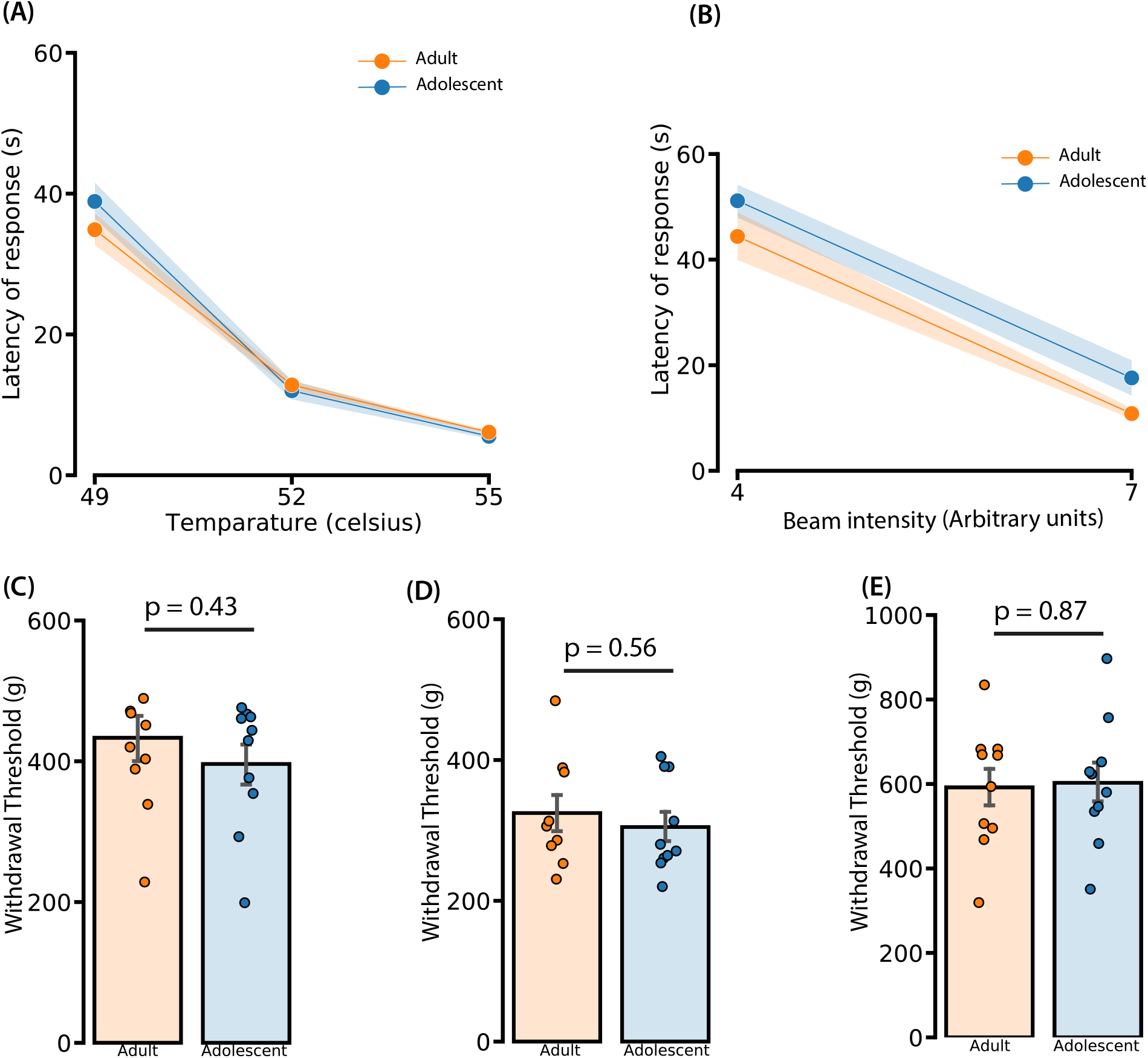
Adolescent (n=12) and adult (n=12) manifested same nociceptive thresholds. **[A] Hot plate test:** A decreased thermal nociceptive threshold for both groups was found either while increasing temperature of the hot plate (Two way repeated measures ANOVA, temperature intensity effect F(2,36) = 207.4, p <0.0001). There was no difference either between groups (F(1,16)=0.43, p=0.51) or an interaction between group and temperature intensity (F(2,36)=1.43, p=0.25). **[B] Tail flick test:** A decreased thermal nociceptive threshold for both groups was found either while increasing intensity of the beam in the tail flick test (Two way repeated measures ANOVA, beam intensity effect F(1,18) = 154.1, p <0.0001). However, there was no difference either between groups (F(1,18)=3.02, p=0.1) or an interaction between group and temperature intensity. **[C**,**D**,**E] Pinch test:** No differences in mechanical sensitivity were found between adolescent and adult rats in the pinch test for (C) left paw (Unpaired T test, t(18)=0.8, p=0.43), (D) right paw (Unpaired T test, t(17)=0.59, p=0.56) and (E) tail (Unpaired T test, t(18)=0.16, p=0.87).

**Supplementary figure 2:**
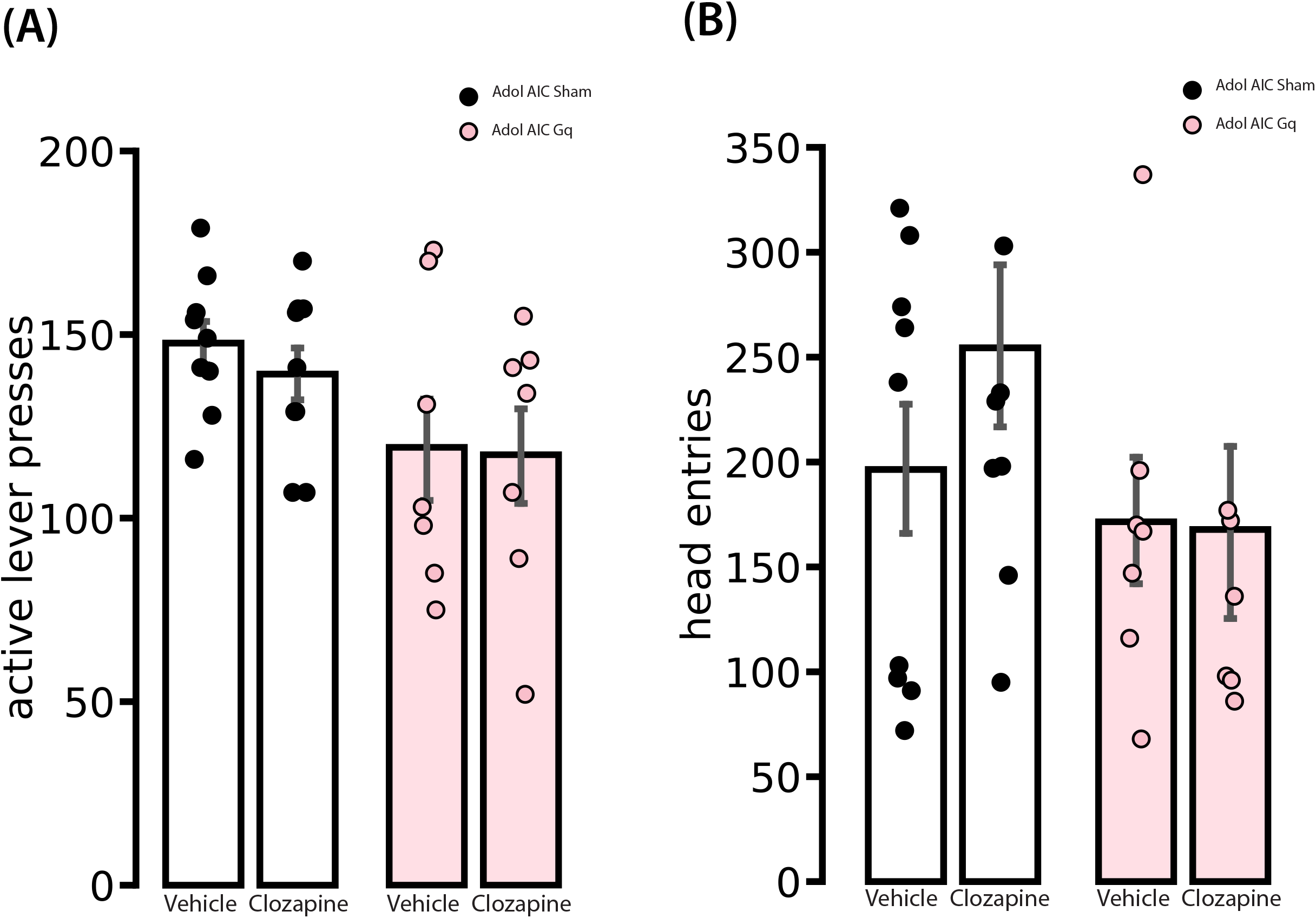
No effects on the motor behavior of adolescent rats due to clozapine. **[A]** On PND 45-46 (see Figure 6A), while rats were trained for baseline operant conditioning (saccharine 0.2%, no foot shock delivery), they received clozapine injection (0.1 mg/kg ip) in a latin square design. As reported on Supplementary Figure 2A, the number of lever presses performed by adolescents rats receiving infusion of the h3MD virus in the AIC was similar to that of rats with sham surgery (Two way repeated measures ANOVA, no group effect F(1,14) = 4.14, p>0.05, no clozapine effect F(1,14) = 0.38, p>0.05, and no interaction effect F(1,14) = 0.14, p>0.05). **[B]** The number of head entries by adolescents rats receiving infusion of the DREADD virus in the AIC as compared to those receiving sham surgery was similar (Two way repeated measures ANOVA, group effect F(1,14) = 1.2, p>0.05) when they were injected with either clozapine or vehicle (Two way repeated measures ANOVA, clozapine effect F(1,14) = 2.02, p>0.05, interaction effect F(1,14) = 2.61, p>0.05) during the compulsivity task.

